# mTOR Inhibition Overcomes RSK3-mediated Resistance to BET Inhibitors in Small Cell Lung Cancer

**DOI:** 10.1101/2021.11.08.467833

**Authors:** Anju Kumari, Lisa Gesumaria, Yan-Jin Liu, V. Keith Hughitt, Xiaohu Zhang, Michele Ceribelli, Kelli M. Wilson, Carleen Klumpp-Thomas, Lu Chen, Crystal McKnight, Zina Itkin, Craig J. Thomas, Beverly A. Mock, David S. Schrump, Haobin Chen

**Author notes:** **Corresponding author**: Haobin Chen, 10 Center Drive Room 3-5888, National Institutes of Health, Bethesda, MD 20892. Senior authors (Craig J. Thomas, Beverly A. Mock, David S. Schrump).

## Abstract

**Purpose:** SCLC is a recalcitrant malignancy with limited treatment options. BET inhibitors have shown promising preclinical activity in SCLC, but their broad sensitivity spectrum limits their clinical prospects in this malignancy. Drug combination could be a solution.

**Experimental design:** We performed high-throughput drug combination screens in SCLC cell lines to identify potential therapeutics synergizing with BET inhibitors. Validation was performed in SCLC cell lines and patient-derived xenograft models. Genome-wide RNA sequencing of xenograft tumors was performed to determine the mechanism underlying the synergy of the drug combination.

**Results:** Inhibitors of the PI-3K-AKT-mTOR pathway were the top candidates from the screens. Among the therapeutics targeting this pathway, mTOR inhibitors showed the highest degree of synergy with BET inhibitors *in vitro*. Furthermore, the combination of these two classes of drugs showed superior antitumor efficacy and tolerability *in vivo*. Using both *in vitro* and *in vivo* SCLC models, we demonstrate that BET inhibitors activate the intrinsic apoptotic cascade, and mTOR inhibitors further enhance these apoptotic effects. Mechanistically, BET inhibitors activate the TSC2-mTOR-p70S6K1 signaling cascade by upregulating RSK3, an upstream kinase of TSC2. Activation of p70S6K1 leads to BAD phosphorylation and cell survival. mTOR inhibition blocks this survival signaling cascade and potentiates the antitumor effects of BET inhibitors.

**Conclusions:** Our results demonstrate that RSK3 upregulation is a novel resistance mechanism of BET inhibitors in SCLC, and mTOR inhibition can overcome this resistance and enhance apoptosis. These findings provide a rationale to evaluate the combination of mTOR and BET inhibitors in patients with SCLC.

## Introduction

Small cell lung cancer (SCLC) is a recalcitrant malignancy with limited treatment options (1). Recent analysis of the Surveillance, Epidemiology, and End Results data found no improvement in the 2-year survival rates of this malignancy from 2001 to 2014 because of limited treatment advances (2). Several new therapeutics, including immune checkpoint inhibitors and lurbinectedin, have been recently approved by the FDA for SCLC, but only a subset of patients receives benefits (3). Therefore, there is an unmet need to develop novel therapies for SCLC.

A potential therapeutic candidate for SCLC is BET inhibitor (BETi). This class of drugs targets the bromodomain and extra-terminal domain (BET) family proteins, namely BRD2, BRD3, BRD4, and BRDT. The primary function of the BET family proteins is gene transcription regulation, and the two bromodomains in each of these proteins are critical for this function by enabling access to active chromatin. BETi binds to these bromodomains and dissociates the BET family proteins from active chromatin, resulting in the suppression of gene transcription. Because BETi only suppresses a subset of genes - particularly those related to cell lineage and driver oncogenes (4), there has been considerable interest in applying this class of drugs for cancer treatment.

A study in preclinical mouse models suggested that SCLC cells are exquisitely susceptible to BETi (5), but a subsequent investigation revealed a broad sensitivity range among human SCLC lines (6). There have been several efforts in developing drug combinations to enhance the antitumoral effects of BETi and to delay the onset of drug resistance. For instance, inhibitors targeting PARP, HDAC6, or BCL2 were found to synergize with BETi in SCLC (7–10). However, to our knowledge, an unbiased screen among the existing oncological therapeutics in combination with BETi has not been reported in SCLC. In addition, the mechanisms underlying BETi resistance are not fully understood in this malignancy.

The PI-3K-AKT-mTOR pathway is commonly activated in cancer, and a recent study identified mTOR as an essential kinase in a subset of SCLC through shRNA library screens (11). Currently, everolimus (an mTOR inhibitor [mTORi]) has been approved for the treatment of multiple neoplasms. However, a phase II trial of everolimus in an unselected patient population of relapsed SCLC only found limited single-agent antitumor activity, suggesting a need to combine this drug with other therapeutics (12). Interestingly, in breast and prostate cancer cell lines, inhibition of mTOR complex 1 (mTORC1) caused a feedback activation of receptor tyrosine kinases (RTKs), which led to phosphorylation at AKT T308 and activation of its downstream signaling to promote survival (13,14). BETi was reported to potentiate the antitumor effects of mTORi by blocking the upregulation of RTKs and thus decreasing this feedback activation in breast and colon cancer cell lines (15,16). To our knowledge, the combination of mTORi and BETi has not been studied in SCLC.

In this study, we performed high-throughput drug combination screens to identify potential therapeutics synergizing with BETi in SCLC in an unbiased manner.

## Materials and Methods

### Cell Culture

H446, H187, H460, H1048, H1436, Calu-6, MDA-MB-468, and HEK293T cells were purchased from the American Type Culture Collection, SCLC-21H was from the German Collection of Microorganisms and Cell Cultures, and COR-L279 and COR-L88 were from Sigma. LX95 SCLC line was established by growing the single cells isolated from the LX95 PDX tumors in HITES medium (DMEM:F12 medium supplemented with 0.005mg/ml insulin, 0.01 mg/ml transferrin, 30 nM sodium selenite, 10 nM hydrocortisone, 10 nM beta-estradiol, 1x Glutamax, 5% heat-inactivated fetal bovine serum, 1x penicillin/streptomycin). H446 cells with ectopic expression of BCL2 were established by transfection of H446 cells with Flag-BCL2 (a gift from Clark Distelhorst (17); Addgene plasmid #18003) followed by selection with 0.4 mg/ml G418 (Roche). All cell lines were grown in culture media recommended by the suppliers and were maintained in humidified incubators at 37°C. MDA-MB-468 was grown at 100% air, while all other lines were at 5% CO_2_. All cell lines tested negative for mycoplasma, and short tandem repeat analysis was performed to authenticate commercial cell lines.

### Reagents

For *in vitro* studies, (±)-JQ1 (Sigma, #SML0974), NHWD870 (a gift from Nenghui Wang and Mingzhu Yin, Ningbo Wenda Pharma), actinomycin D (Sigma, #A9415), everolimus (Sellekchem, #S1120), LJH685 (Sellekchem, #S7870), MK-2206 (Sellekchem, #S1078), PF-4708671 (Sellekchem, #S2163), rapamycin (Sellekchem, #S1039), torkinib (Sellekchem, #S2218), and Z-VAD-FMK (Sigma, # 219007) were dissolved in DMSO; copanlisib (Sellekchem, #S2802) was prepared in 10% trifluoroacetic acid (Sigma, #T6508).

For *in vivo* studies, AZD5153 (Chemitek, #CT-A5153), AZD5363 (Chemitek, #CT-A5363) and NHWD870 were first dissolved in N,N-dimethylacetamide (Sigma, #270555) and then mixed with 0.5% methylcellulose (Sigma, #M0430) plus 0.1% Tween-80 (Sigma, #P4780) before administration. Everolimus was first dissolved in propylene glycol (Sigma, # W294004) at 10mg/600μl concentration and was then combined with 1ml 10% Tween 80, and finally with 400μl water. After mixing, the drug solution was mixed with 0.5% methylcellulose plus 0.1% Tween-80 prior to each dosing.

### High-Throughput Combinatorial Screens

High-throughput combinatorial screens were performed as previously described (18) with some modifications. The initial screen was performed in H446 cells by combining JQ1 with each drug from a Mechanism Interrogation PlatE (MIPE) 4.0 library in a 6×6 matrix layout. The MIPE 4.0 library consists of 1,912 FDA-approved oncological therapeutics and investigational agents. The viability endpoint of the initial screen was measured using the CellTiterGlo® reagent (Promega) 72 hours after drug treatment. The synergy of various drug combinations was assessed using the excess Highest Single Agent (HSA) method, the response heat map, and the ΔBliss heat map (19,20).

The second screen was performed in four SCLC cell lines (H446, COR-L279, H187, and SCLC-21H) in a 10×10 matrix by combining JQ1 or iBET-762 with one of the 40 drugs selected from the first screen. The endpoints of the second screen were caspase 3/7 activity measured using the CaspaseGlo^®^ 3/7 assay reagent (Promega) at the 8-hour and 16-hour time intervals, as well as cell viability measured using the CellTiterGlo^®^ reagent at the 72-hour time interval following drug treatment.

### Cell Viability and Caspase 3/7 Activity Assay

Cells were dissociated with TrypLE (ThermoFisher), and large cell clumps were removed by passing the cell solution through sterile 40 μm nylon mesh cell strainers (Fisher Scientific, Hampton, NH). Subsequently, 750 cells in 15 μl growth media were seeded into each well of a 384-well plate (Corning, #3765). On the next day, drugs were serially diluted 2-fold in appropriate solvents to generate nine consecutive concentrations. Then, the compound and its vehicle control were diluted 100 times in culture media and delivered to the cells in 10% of the final volume to achieve a total of 1,000X dilution. Assays were performed with four replicates for each dose. Caspase 3/7 activity was measured using the caspase-glo^®^ 3/7 Assay system (Promega) 48 hours after drug treatment, and cell viability was examined at the 72-hour interval using the CellTiter-Glo^®^ 2.0 Reagent (Promega). A reflective foil seal (Bio-Rad, #MSF1001) was applied to the bottom of each plate to maximize the output signal.

### Transfection

Lipofectamine 2000 or 3000 reagents (Thermofisher) were used for lipofection by following the manufacturer’s recommendations. The Myr-RSK3 vector (pWZL Neo Myr Flag RPS6KA2) was a kind gift from William Hahn & Jean Zhao (Addgene plasmid # 20621; (21)).

### Western Blot Analysis

After cells were collected and washed with PBS, cells were lysed in cold 1X RIPA lysis buffer (Millipore) supplemented with a protease inhibitor cocktail (Sigma). When a phosphorylated protein was measured, RIPA lysis buffer was also complemented with a phosphatase inhibitor cocktail (Sigma or ThermoFisher). After incubation on ice for 10 minutes, cells were centrifuged at 10,000 *g* for 10 minutes to extract cell lysates. Mitochondrial fractions (for cytochrome c measurement) and membrane fractions (for Myr-RSK3 measurement) were extracted from cells using a Cell Fractionation kit (Abcam) and a Mem-PER™ Plus Membrane Protein Extraction Kit (Thermofisher), respectively.

Proteins were quantified using a Pierce BCA Protein Assay Kit (ThermoFisher). Equivalent amounts of protein were resolved on precast polyacrylamide denaturing gels and were then transferred onto 0.2 μm PVDF membranes using either dry transfer with an iBLOT2 system (ThermoFisher) or conventional wet transfer. Membranes were incubated with primary antibodies at the concentrations specified in Table S1. After membrane incubation with an appropriate secondary antibody, the presence of a protein of interest was detected by chemical fluorescence following a conventional ECL Western blotting protocol. Densitometry analysis was performed using the Image Lab software (Biorad).

### Animal Studies

The animal experiments were approved and performed according to the regulations set by the National Cancer Institute-Bethesda Animal Care and Use Committee. SCLC patient-derived xenograft (PDX) LX33 and LX95 models were generously provided by Charles M. Rudin and John T. Poirier (22,23). Freshly isolated PDX tumors were dissociated into single-cell suspensions using the Human Tumor Dissociation Kit (Miltenyi Biotec) and a gentleMACS™ Octo Dissociator (Miltenyi Biotec) following the manufacturer’s instructions. After washing cells with PBS three times, red blood cells were lysed by resuspending cells in 5 ml ACK lysis buffer (Quality Biological) followed by incubation at room temperature for 5 minutes. After another three washes with PBS, approximately 5×10^6^ viable cells in 100 μl PBS were injected subcutaneously into the right flanks of 6-week-old NOD-SCID mice (Charles River Laboratories).

Once the tumor volume reached 50-100 mm^3^ (average, 14-21 days), mice were randomized into each treatment group and were given vehicle, NHWD870 or AZD5153, everolimus or AZD5363, or a combination of two drugs via oral gavage. The dosages and frequencies of drug administration are specified in the figure legends. Body weights and tumor sizes were measured every three days. Treatment was on hold if one of the following two criteria was met: 1) >15% body weight loss compared to the initial body weight (at the point of randomization); or 2) active diarrhea. Treatment was resumed once the bodyweight loss recovered to <10% reduction of the initial weight and diarrhea had stopped for three days. Animals were euthanized if 1) tumor volume was ≥1500 mm^3^; 2) tumor became ulcerated; 3) 75 days had elapsed after a tumor became palpable.

### Histology Analysis

Formalin-fixed tumors were embedded in paraffin wax, and sectioned tissues were stained with hematoxylin and eosin or with anti-cleaved caspase 3 (Cell Signaling Technology, #4850, 1:200) and anti-RSK3 antibodies (Novus Biologicals, NBP2-52555, 1:200). Quantification of cleaved caspase 3-positive cells was performed using QuPath [v0.2.2; (24)] by counting cytoplasmic staining-positive cells in 4-15 randomly selected images of each tumor.

### RNA Isolation from Tumor Cell of PDX

Tumor cells were isolated from PDX using a magnetic-activated cell sorting method (25). In brief, after obtaining single-cell suspensions from fresh tumors, cells were centrifuged and resuspended at concentrations of 2×10^6^ cells per 80 μl PBS supplemented with 0.5% bovine serum albumin (Sigma). Afterward, cells were mixed with a 20μl cocktail from a Mouse Cell Depletion Kit (Miltenyi Biotec). Separation of mouse cells from human tumor cells was achieved using an LS column on a MACS Separator (Miltenyi Biotec). The flow-through fraction, which contained enriched human tumor cells, was collected by centrifugation. The cell pellet was processed for total RNA isolation using an RNeasy Mini Kit (Qiagen).

### RNA Sequencing (RNA-seq)

mRNA libraries were constructed using a TruSeq Stranded mRNA Library Prep kit V2 (Illumina) and were paired-end sequenced on a HiSeq 4000 system (Illumina) at the NCI Frederick Sequencing core facility.

RNA-seq reads were processed using the Snakemake-based lcdb-wf transcriptomics pipeline v1.3c (https://github.com/lcdb/lcdb-wf) (26). Briefly, quality was assessed using FastQ Screen and FastQC, and adapter sequences and low-quality base pairs were trimmed using cutadapt. Trimmed reads were then mapped to the human reference genome (gencode v30) using HISAT2 (v2.1.0), and transcript levels were quantified using the *featureCounts* method from RSubRead (v1.6.4). Differential expression analysis was performed in the R/Bioconductor environment (R v4.0.2; Bioconductor v2.48.0) using the DESeq2 package (v1.28.0) (27,28). After constructing a DESeqDataSet object from the read counts generated by RSubRead, generalized linear models were fit with default parameters using a treatment coding design (e.g., NHWD870 vs. Vehicle). Finally, shrunken log-fold changes were estimated for each contrast using the *apeglm* adaptive t prior shrinkage approach (29).

### Quantitative Reverse-transcription PCR (RT-PCR) Assay

After total RNA isolation, RNA was quantified using Nanodrop. Equivalent amounts of total RNA were used for cDNA synthesis using an iScript cDNA Synthesis kit (Bio-Rad). Quantitative analysis of *RPS6KA2* was performed using a SsoAdvanced™ Universal SYBR^®^ Green Supermix (Biorad) and a set of specific primers (*RPS6KA2*, Biorad, qHsaCED0042305; β-actin, Biorad, qHsaCED0036269) on an ABI PRISM7500 real-time PCR system (ThermoFisher).

### Data Analysis and Statistical Analysis

GraphPad Prism 8.0 was used for statistical analysis and graphing. CompuSyn (www.combosyn.com) was used to determine drug synergy of the low-throughput experiments by calculating combination indexes (30). Two-sample comparisons were performed using the Student’s unpaired *t*-test, and multiple sample comparisons were analyzed using the ANOVA test with multiple comparison correction (Dunnett’s method). The Log-rank (Mantel-Cox) test was employed for Kaplan-Meier survival analyses. A two-tailed p<0.05 was considered statistically significant. Error bars represent mean ± SD.

### Data Availability

All screen results are publicly accessible at https://tripod.nih.gov/matrix-client/. RNA-seq raw data were deposited in the GEO database (GSE155923).

## Results

### mTORi synergizes with BETi *in vitro*

We first examined the combination of JQ1 (a prototype BETi) and a MIPE 4.0 library of 1,912 approved and investigational drugs, using a 6×6 matrix layout and an endpoint of cell viability. The outcomes of this experiment demonstrated a strong enrichment for mTORi agents (e.g., rapamycin, everolimus, ridaforolimus, and temsirolimus) among the highest-ranked synergetic drug-drug pairs (Fig. 1a and Table S2).

**Figure 1.**
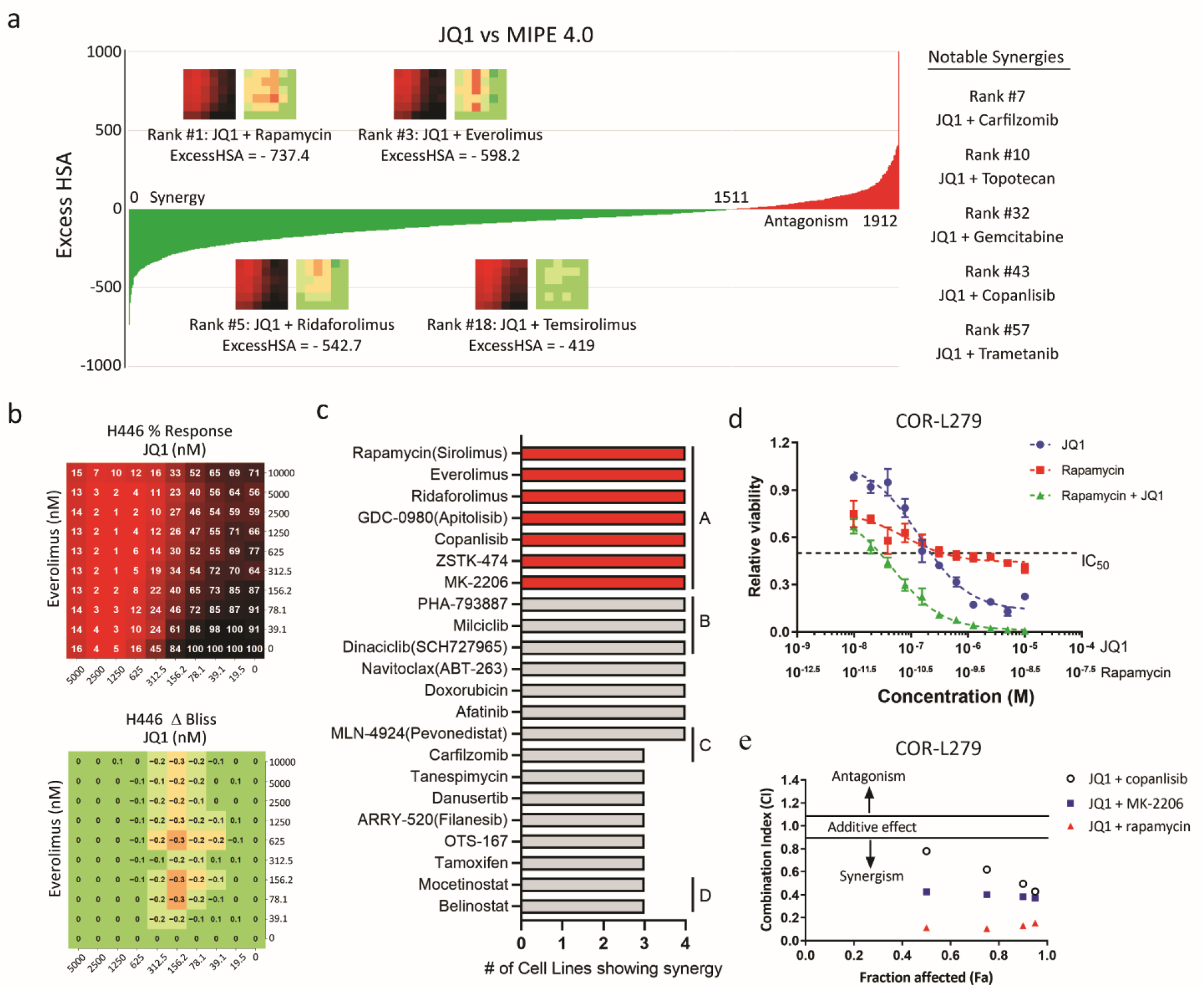
mTORi synergizes with BETi *in vitro.* **(a)** Ranking of synergy and antagonism of JQ1 in combination with 1,912 agents in the MIPE (Mechanism of Interrogation Plate) 4.0 library, according to the Excess HSA (Highest Single Agency) metric in the primary screens in H446 cells. Prominent drug synergies are shown, including the combination of JQ1 with several mTORi agents. **(b)** The percentage response (top) and Δ Bliss (bottom) heatmaps for the 10 × 10 screen of the JQ1 and everolimus combination in H446 cells. **(c)** The candidates that synergized with BETi in at least three SCLC lines in the 10 × 10 screens. A, B, C, and D represent inhibitors targeting the PI-3K-AKT-mTOR pathway, cyclin-dependent kinases, proteolysis, and HDAC, respectively. **(d)** Relative viability of COR-L279 cells at the 72-hour time interval after treatment of JQ1, rapamycin, or their combination. **(e)** Synergy plot showing the combination index (CI) versus affected fractions calculated based on the viability results of the specified drug-drug combinations in COR-L279 cells at the 72-hour time interval.

Follow-up studies explored 40 of the most promising outcomes using an expanded 10×10 matrix layout with optimized concentration ranges. JQ1 and another BETi, I-BET762, were tested in parallel across 4 SCLC cell lines (H446, COR-L279, H187, and SCLC-21H). Besides cell viability at 72 hours after treatment, we also measured apoptosis induction at the 8- and 16-hour time intervals. Figure 1b shows a typical example of drug synergy heatmaps. We identified 22 drugs synergistic with JQ1 and I-BET762 in 3 or 4 out of the 4 SCLC cell lines, with inhibitors targeting the PI-3K-AKT-mTOR pathway being on the top of the list (Fig. 1c).

To validate the results of the high-throughput screens, we performed independent viability assays using a fixed concentration ratio of two-drug combinations. As shown in one typical example in Figure 1d, concurrent treatment with rapamycin decreased the IC_50_ value of JQ1 by 10-fold in COR-L279 cells. We quantified the synergy between JQ1 and the inhibitors targeting PI-3K (copanlisib), AKT (MK-2206), or mTOR (rapamycin) using the method of Chou and Talalay (30). On the basis of the combination index (CI) at four different dose levels that caused a 50%, 75%, 90%, and 95% decrease in cell viability (fraction affected), the combination of JQ1 and rapamycin showed a higher degree of synergy than the other two combinations in COR-L279, H446, and H187 cells (Figs. 1e and S1a). Because the toxicity profiles of mTORi and BETi are non-overlapping and their combination shows the highest degree of synergy, we focused on this combination for subsequent study.

To ensure the synergy between rapamycin and JQ1 was not due to an off-target effect, we evaluated the combinations of JQ1 and various mTORi agents (first generation: rapamycin and everolimus; second-generation: torkinib), as well as the cotreatment of everolimus and a different BETi (NHWD870; (31)). As shown in Figure S1b, the combinations of various mTOR and BET inhibitors consistently demonstrated a strong synergy. We also tested the combination of JQ1 and everolimus in SCLC cell lines resistant to JQ1 (IC_50_ ≥ 20μM), and a strong synergy was observed in 2 out of 3 cell lines (Fig. S1c). Collectively, these results demonstrate that mTORi potentiates the growth inhibitory effects of BETi in SCLC lines.

### mTOR inhibition amplifies BETi-induced apoptosis in SCLC

We next asked whether the combinations of mTORi and BETi induce apoptosis in SCLC. To answer this, we treated COR-L279 cells with the combination of JQ1 and rapamycin in the presence or absence of a pan-caspase inhibitor, Z-VAD-FMK. As shown in Figure 2a, Z-VAD-FMK drastically attenuated the decrease in cell viability caused by the JQ1 and rapamycin combo, suggesting that apoptosis is partly responsible for growth inhibition. Apoptosis can be triggered through either the intrinsic or extrinsic cascade, with the former one characterized by cleavage of caspase 9 and the latter by cleavage of caspase 8 (Fig. 2b). Both pathways converge on the cleavage of caspase 3 and PARP to activate cell death, and crosstalk between these two pathways can occur through cleaved BID (also known as t-BID).

**Figure 2.**
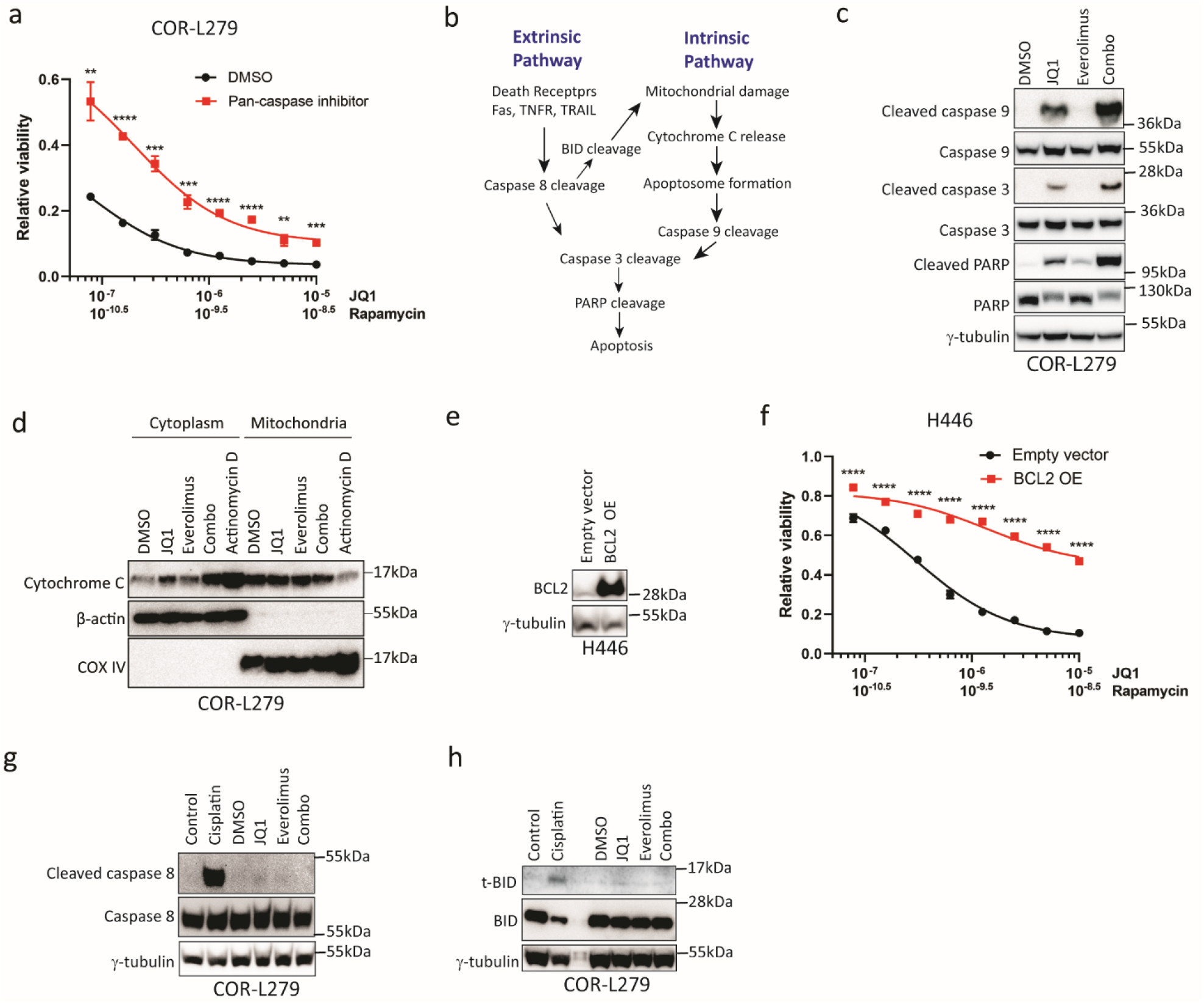
mTORi amplifies BETi-induced apoptosis in SCLC via the intrinsic apoptotic cascade. **(a)** Relative viability of COR-L279 cells at 72 hours after treatment with the JQ1 and rapamycin combo with or without a pan-caspase inhibitor, Z-VAD-FMK (2.5μM). **(b)** A diagram illustrating the intrinsic and extrinsic apoptotic cascades. **(c)** Western blots show caspase 9, caspase 3, and PARP cleavage in COR-L279 cells at 24 hours following treatment of JQ1 (1μM), everolimus (6.25nM), or their combination. **(d)** Western blots show cytochrome c abundance in mitochondria and cytoplasmic fractions of COR-L279 cells, at 16 hours after treatment of JQ1 (1μM), everolimus (6.25nM), or their combination. Actinomycin D (1μg/ml, 16 hours) was used as a positive control. β-actin and Cox IV serve as loading controls for the cytoplasmic and mitochondrial fractions, respectively. **(e)** Ectopic expression of BCL2 in H446 cells as assessed by immunoblotting. **(f)** Relative viability of H446 cells with ectopic expression of BCL2 versus control cells at 72 hours after treatment of JQ1, rapamycin, or the combination. **(g-h)** Western blots show cleaved caspase 8 (g) and t-BID (h) in COR-L279 cells following treatment of JQ1 (1μM), everolimus (6.25nM), or the combination. Cisplatin at 20μM and 50μM were used as positive controls in (g) and (h), respectively. The significance of the two-group comparisons was determined using the Student’s *t*-test in (a) and (f). Error bars represent SD; **, p<0.01; ***, p<0.001; ****, P<0.0001. OE, overexpression.

To dissect which apoptotic pathway is activated by the mTORi and BETi combo, we first examined cleaved caspase 9, a key intermediate of the intrinsic cascade. As shown in Figure 2c, JQ1 increased cleaved caspase 9 and the downstream cleaved caspase 3 and PARP; the addition of everolimus further increased these cleaved apoptotic proteins, suggesting that JQ1 and the combo activated the intrinsic apoptotic pathway. Because the intrinsic apoptotic cascade is typically triggered by the release of cytochrome c from mitochondria (Fig. 2b), we isolated cytosolic and mitochondrial fractions of COR-L279 cells after drug treatments. As shown in Figure 2d, cytochrome c was elevated in the cytosolic fraction of the JQ1-treated cells and was substantially higher in the sample treated with the everolimus and JQ1 combination, providing further evidence of involvement of the intrinsic apoptotic pathway. To further confirm that the intrinsic pathway is activated, we utilized an H446 cell line stably expressing ectopic BCL2, an antiapoptotic protein that stabilizes mitochondrial membrane and blocks the efflux of cytochrome c (Fig. 2e). The JQ1 and rapamycin combination was less effective in suppressing the growth of cells with ectopic expression of BCL2 relative to vector control, confirming that the intrinsic apoptotic pathway was activated by the drug combination (Fig. 2f).

We next assessed whether the everolimus and JQ1 combination activates the extrinsic apoptotic pathway. As shown in Figure 2g, cisplatin, but not JQ1 or its combination with everolimus, induced cleaved caspase 8 in COR-L279 cells. As expected, t-BID was elevated in cells treated by cisplatin but not in those that received JQ1 or the drug combo (Fig. 2h). Taken together, these results demonstrate that the mTORi and BETi combination induces apoptosis by activating the intrinsic apoptotic pathway in SCLC.

### mTOR inhibition enhances the antitumor effects of BETi *in vivo* by increasing apoptosis

To evaluate the antitumor effects of the mTORi and BETi combo *in vivo*, we tested the combination of everolimus and NHWD870 in the SCLC PDX model LX33. As shown in Figure 3a-b, compared to either single drug, concurrent administration of everolimus and NHWD870 resulted in significantly better control of tumor growth and a longer median overall survival (50 days in the combo group versus 30 days in single-agent groups). In another SCLC PDX model (LX95), we tested the combination of everolimus and AZD5153, a bivalent BETi in phase I/II clinical trials (32). Like the combination of everolimus and NHWD870 in LX33, concurrent administration of everolimus and AZD5153 achieved significantly better control of tumor growth and increased the median overall survival from 30 days in single-drug groups to 60 days (Fig. 3c-d). Both drug combinations were reasonably tolerated, except for mild diarrhea or weight loss in a few animals. These side effects commonly improved spontaneously after holding the treatments for a few days.

**Figure 3.**
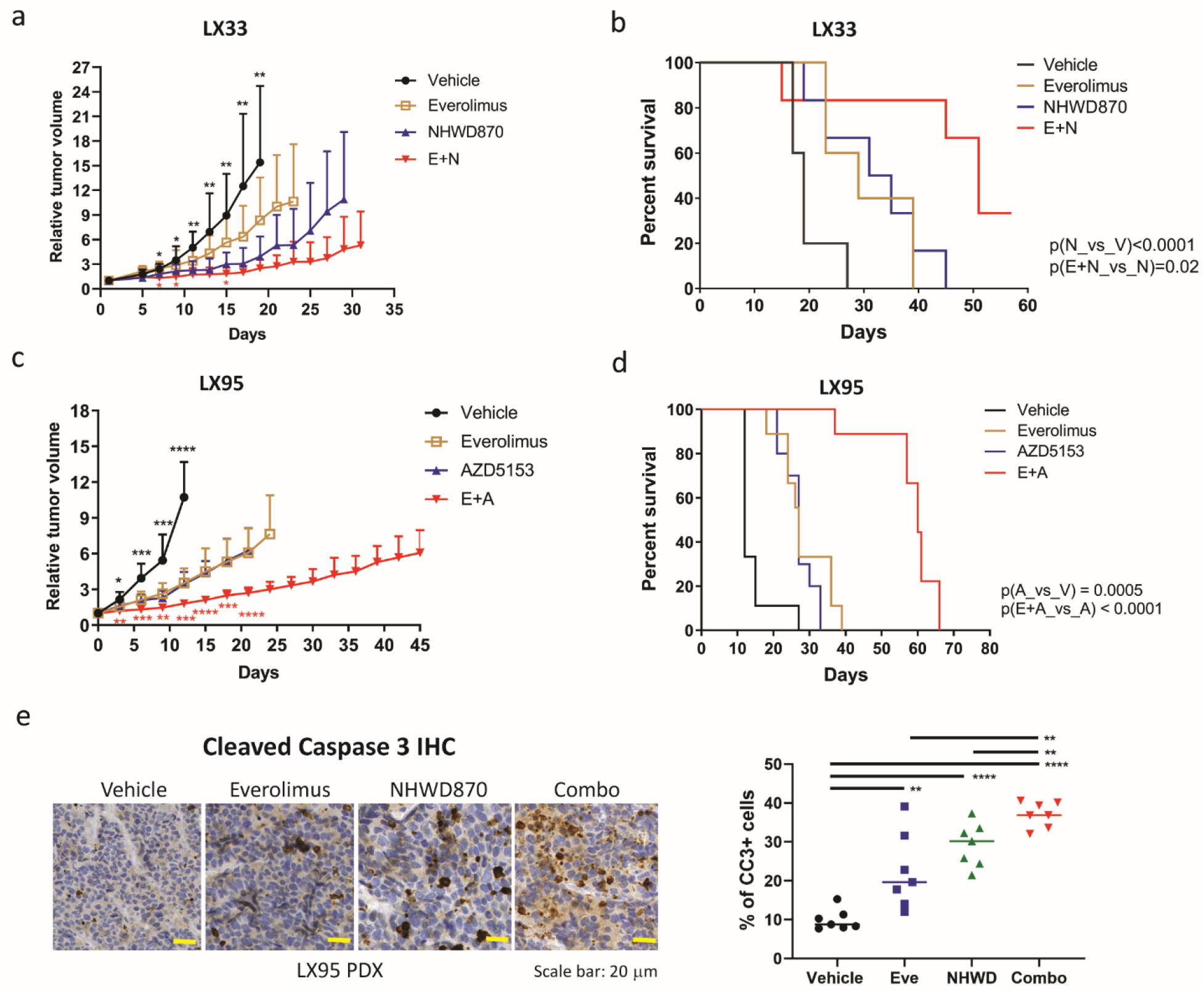
mTOR inhibition enhances the antitumor effects of BETi *in vivo* by increasing apoptosis. **(a-b)** Compared to either single agent, the combination of everolimus (3mg/kg, daily) and NHWD870 (3mg/kg, daily) was more effective in controlling tumor growth (a) and prolonging survival (b) in the LX33 SCLC PDX model (n ≥ 10 per group). **(c-d)** Compared to single-agent everolimus (2mg/kg, daily) and AZD5153 (1.5 mg/kg, daily), concurrent treatment with both drugs at 1mg/kg daily was more effective in controlling tumor growth (c) and prolonging survival (d) in the LX95 model (n=9 per group). (e) Representative images (left) and quantification (right) of cleaved caspase 3 IHC staining in LX95 PDX tumors after one-week treatment of everolimus (2mg/kg, daily), NHWD870 (1.5mg/kg, daily), or the combination (everolimus 1.5mg/kg and NHWD870 1mg/kg, daily). n=7 per group. Horizontal lines in the right panel represent medians. The significance of the two-group comparisons was determined using the Student’s *t*-test in (a) and and Log-rank test in (b) and (d). Statistically significant differences in tumor volumes between the BETi single-agent and vehicle groups are indicated with black asterisks, while those between the combo and BETi single-agent groups are marked with red asterisks. Error bars represent SD; *, p<0.05; **, p<0.01; ***, p<0.001; ****, p<0.0001. A, AZD5153; E, everolimus; N, NHWD870; V, vehicle; CC3, cleaved caspase 3; IHC, immunohistochemical staining.

We also evaluated the combination of AKT inhibitor (AKTi) and BETi *in vivo*. Compared to single-agent AZD5153, the combination of AZD5363 (AKTi) and AZD5153 did not lead to better control of tumor growth or a longer median overall survival in the LX95 model (Fig. S2a-b). The lack of improvement was due to excessive toxicity, as this combination caused severe weight loss, diarrhea, and premature death, none of which occurred in single-agent groups. Despite a 40% dose-reduction of AZD5363, this combination was still difficult to tolerate and necessitated extended treatment delays.

We next assessed whether the combination of mTORi and BETi induced apoptosis *in vivo*. To this end, we measured the expression of cleaved caspase 3 in LX95 tumors after one-week treatment of everolimus and NHWD870. Consistent with our *in vitro* findings, tumor cells positive for cleaved caspase 3 were more commonly present in the xenografts from the drug combo group relative to those in the single-agent groups (Fig. 3e). We next used a tumor cell line established from the LX95 PDX to evaluate whether the drug treatments activated the intrinsic apoptotic pathway. We first confirmed this cell line formed similar xenograft tumors as those grown from the implanted LX95 tumor blocks (Fig. S3a). Like COR-L279, the combination of everolimus and JQ1 activated the intrinsic apoptotic pathway by increasing cleavage of caspase 9 and release of cytochrome c from mitochondria (Fig. S3b-c). We next examined the role of the extrinsic apoptotic pathway. In agreement with a previous report that this gene is frequently lost in SCLC (33), caspase 8 was barely detectable in LX95 cells (Fig. S3d). Taken together, these results demonstrate that mTORi, but not AKTi, potentiates the antitumor effects of BETi *in vivo* by increasing apoptosis via the intrinsic apoptotic cascade.

### BETi upregulates RSK3, a potential upstream kinase of the TSC2-mTOR cascade, in SCLC

It was previously reported that BETi synergizes with mTORi in breast and colon cancer cell lines by blocking the mTORi-induced RTK feedback activation (15,16). To determine whether this mechanism explains the synergy of the mTORi and BETi combo in SCLC, we compared the changes in AKT T308 phosphorylation following mTOR inhibition between a breast cancer cell line MDA-MB-468 and two SCLC cell lines, COR-L279 and LX95. As shown in Figure S4a-c, a steady increase in AKT T308 phosphorylation following mTOR inhibition, indicative of RTK feedback activation (13–16), was observed only in MDA-MB-468 cells but not in COR-L279 and LX95 cells. These findings suggest that, in SCLC, mTOR inhibition does not induce RTK feedback activation and a different mechanism is responsible for the synergy between mTORi and BETi.

To identify the underlying mechanism of drug synergy, we performed RNA sequencing to profile LX95 PDXs following one-week treatment with everolimus, NHWD870, or their combination. Principal-component analysis showed that the transcriptomes of the tumor cells were clustered into four treatment groups (Fig. 4a). By comparing the NHWD870-treated tumors to control tumors, a total of 929 genes had at least a 2-fold change in expression with an adjusted P-value less than 0.05. Ingenuity Pathway Analysis of these identified genes correctly predicted that BRD4 was inactivated by NHWD870 treatment, while TSC2, an upstream regulator of mTOR signaling, was among the top activated regulators (Fig. 4b). Interestingly, one of the 20 most upregulated genes in the NHWD870 and combo groups was *RPS6KA2* (Fig. 4c), which belongs to the same family of *RPS6KA1 –* a known upstream kinase of TSC2 (34). Using quantitative RT-PCR and immunohistochemical staining, we confirmed that *RPS6KA2* and its encoded protein - RSK3, were induced by NHWD870 or the combo in LX95 tumors (Fig. 4d-e).

**Figure 4.**
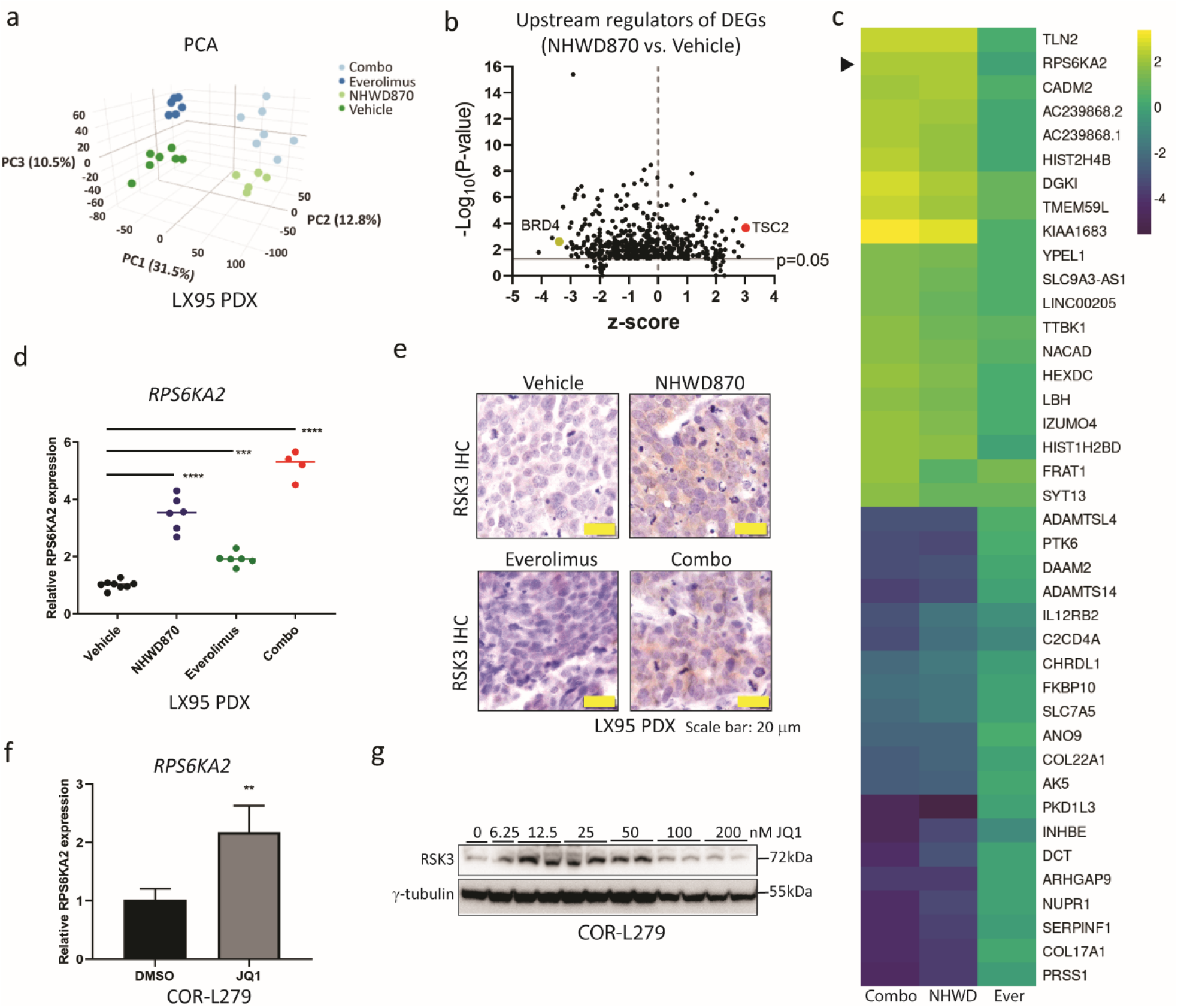
BETi upregulates *RPS6KA2* and its encoded protein RSK3 in SCLC *in vitro* and *in vivo*. **(a)** Principal Component Analysis plot showing clustering of LX95 PDXs treated with everolimus (2mg/kg, daily), NHWD870 (1.5mg/kg, daily), the combination (everolimus 1.5mg/kg and NHWD870 1mg/kg, daily), or vehicle (daily) for one week. **(b)** The volcano plot for the statistical significance (Y-axis) versus the degree of activation or inhibition (X-axis) of the upstream regulators predicted by Ingenuity Pathway Analysis. The input data was the differentially expressed genes (>2 fold-change and adjusted p-value <0.05) comparing the tumors in the NHWD870 group vs. control tumors. **(c)** Heatmap of the top 20 up- and down-regulated genes in the combo group relative to the vehicle group. The arrowhead points to the heatmap of *RPS6KA2*. **(d-e)** qRT-PCR and IHC staining assessing RSK3 gene and protein expression in the LX95 PDX tumors treated with vehicle, NHWD870, everolimus, or the combo. **(f-g)** RSK3 gene and protein expression in COR-L279 cells following one-week treatment of JQ1 at 50nM (f) or the specified doses (g). Fresh media and JQ1 were replenished every three days. The significance of the two-group comparisons was determined using the Student’s *t*-test in (f) and the ANOVA test with multiple comparison correction in (d). Error bars represent SD; **, p<0.01; ***, p<0.001; ****, p<0.0001.

We modeled the *in vivo* experiment by treating COR-L279 cells with various doses of JQ1 for one week *in vitro*. JQ1 was confirmed to upregulate the gene and protein expression of RSK3 in the dose range of 12.5 to 50nM (Fig. 4f-g). Collectively, these results demonstrate that BETi induces RSK3, a potential upstream kinase of TSC2, in SCLC.

### RSK3 induction increases resistance to BETi-induced apoptosis

To evaluate the functional consequence of RSK3 induction, we overexpressed a myristoylated (myr) RSK3 in COR-L279 cells. The addition of a myr sequence recruits overexpressed RSK3 to the cell membrane and subsequent activation of its kinase activity (21). Compared to empty vector control, forced expression of myr-RSK3 increased phosphorylation at TSC2 S1798, demonstrating that TSC2 is a kinase substrate of RSK3 (Fig. 5a). Phosphorylation at p70S6K1 T389, a direct downstream target of mTORC1, was also increased in the RSK3-overexpressed cells, indicating that RSK3 augments mTOR signaling (Fig. 5a). To determine whether BETi-induced RSK3 could activate the TSC2-mTOR-p70S6K1 signaling, we treated COR-L279 cells with JQ1 at 50nM for one week. As shown in Figure 5b, a similar increase in phosphorylation at both TSC2 S1798 and p70S6K1 T389 was observed, demonstrating that BETi activates TSC2-mTOR signaling via RSK3 upregulation.

**Figure 5.**
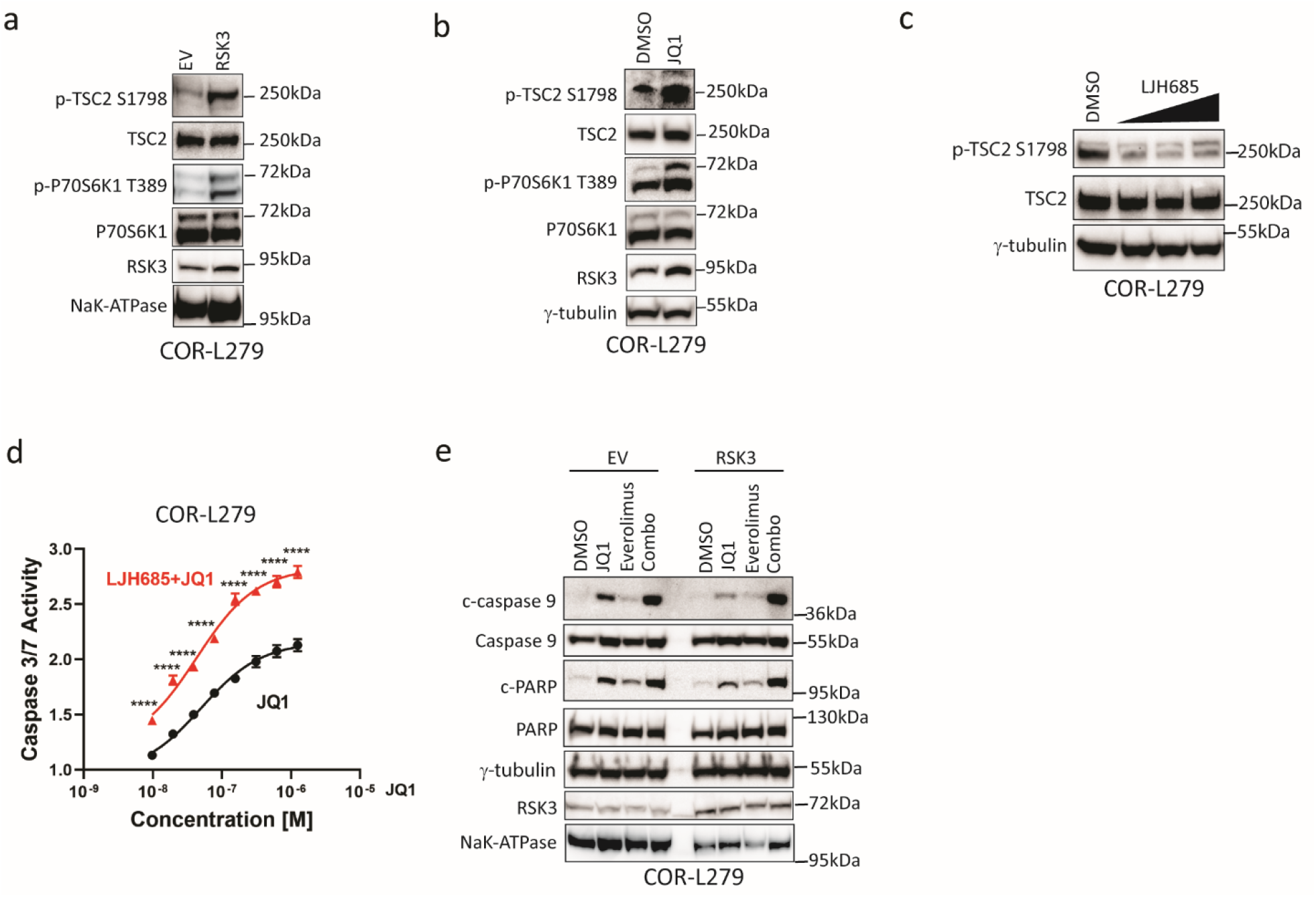
RSK3 upregulation causes resistance to BETi-induced apoptosis. **(a)** Phosphorylation at TSC2 S1798 and p70S6K1 T389 in COR-L279 cells after forced expression of myr-RSK3 versus empty vectors (EV). NaK-ATPase was used as a loading control for membrane fraction proteins. **(b)** Phosphorylation status at TSC2 S1798 and p70S6K1 T389 in COR-L279 cells treated with JQ1 (50 nM, 7 days) versus control cells. **(c)** Reduction of phosphorylation at TSC2 S1798 following 24-hour treatment of LJH685 at 5, 10, 20 μM. **(d)** LJH685 (10 μM) enhanced JQ1-induced caspase 3/7 activation in COR-L279 cells at 48 hours post-treatment. **(e)** Overexpression of myr-RSK3 attenuated apoptosis caused by JQ1 but not by the combo. One day after transfection with Myr-RSK3 expression vector or empty vector (EV), COR-L279 cells were treated with JQ1 (1 μM), everolimus (10 nM), or the combination for 24 hours before cleaved PARP and caspase 9 were assessed by immunoblotting. The significance of the two-group comparisons was determined using the Student’s *t*-test in (d). The error bars represent SD; ****, P<0.0001.

To evaluate how RSK3 upregulation would affect cell response to BETi, we first used LJH685, a selective inhibitor of the RSK family kinases (35), to block TSC2 phosphorylation by RSK kinases (Fig. 5c). Figure 5d shows that LJH685 enhanced caspase 3/7 activation following JQ1 treatment in COR-L279 cells. Next, we overexpressed myr-RSK3 in COR-L279 cells and observed a decrease in JQ1-induced caspase 9 and PARP cleavage, suggesting that RSK3 induction attenuates JQ1-induced apoptosis (Fig. 5e). Everolimus abrogated the protective effect of RSK3 overexpression, as the combo induced the same levels of cleaved caspase 9 and PARP in RSK3-overexpressed cells compared to control cells (Fig. 5e). Collectively, these findings demonstrate that BETi-induced RSK3 activates TSC2-mTOR signaling and causes resistance to BETi-induced apoptosis; and mTORi abolishes this protective effect.

### p70S6K1 mediates the antiapoptotic effects of RSK3 by phosphorylating BAD

To determine whether p70S6K1 mediates the antiapoptotic effects of RSK3, we treated COR-L279 cells with JQ1 with or without PF-4708671, a specific inhibitor of p70S6K1. As shown in Fig. 6a, the combination of PF-4708671 and JQ1 recapitulated the increased apoptosis caused by the mTORi and BETi combination. To further define the role of p70S6K1, we overexpressed a constitutively active form of rat S6K1 (S6K1-deltaCT) and the wildtype (S6K1-WT) in COR-L279 cells. As shown in Figure 6b-c, S6K1-deltaCT, but not S6K1-WT, increased the viability and decreased apoptosis after JQ1 treatment, confirming that p70S6K1 mediates the antiapoptotic effects of RSK3.

**Figure 6.**
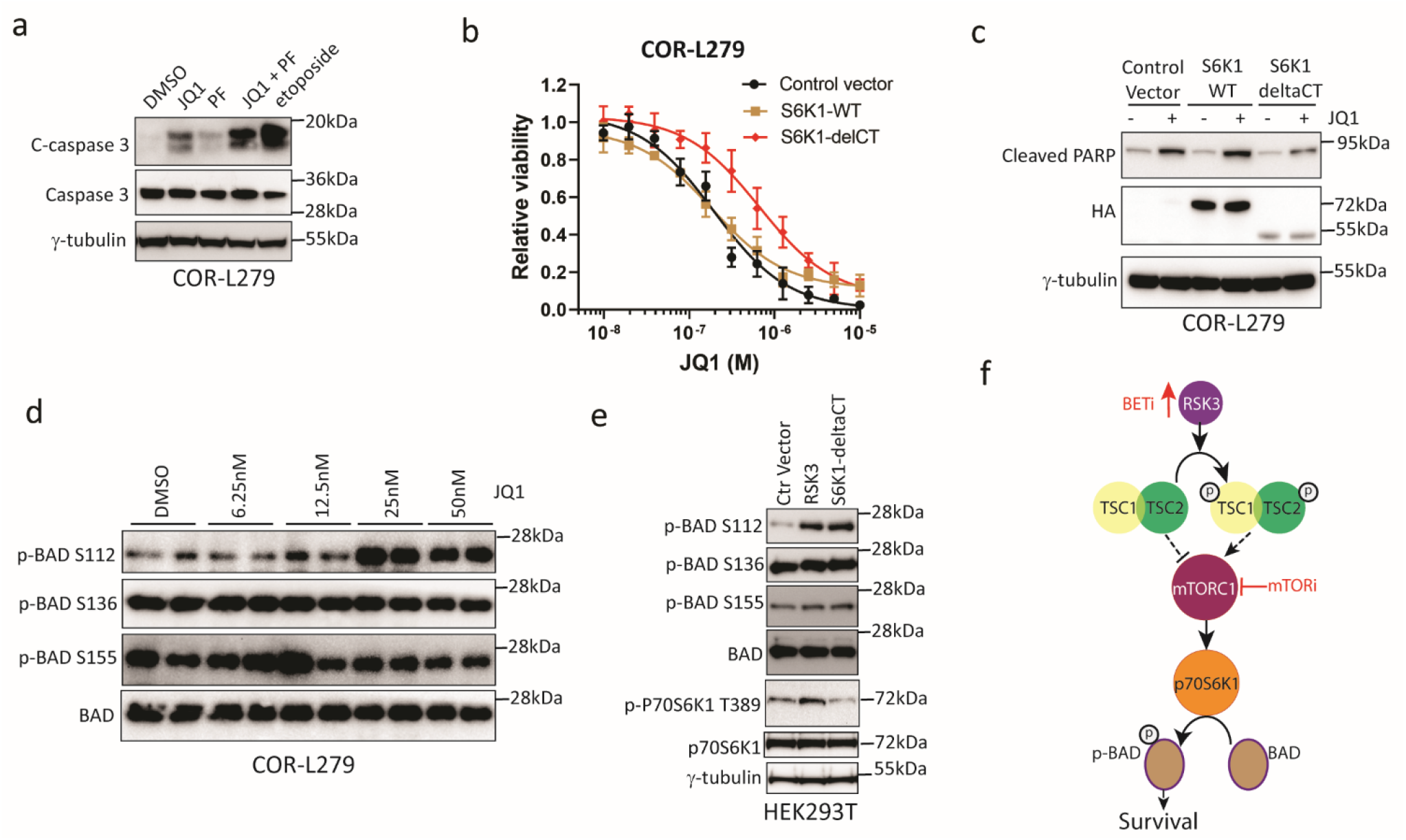
p70S6K1 mediates the antiapoptotic effects of BETi-induced RSK3. **(a)** Cleavage of caspase 3 in COR-L279 cells at 24 hours after treatment of JQ1 (1.67μM), PF-4708671 (PF; 16μM), their combination. Etoposide serves as a positive control. **(b-c)** Overexpression of an active form of S6K1 (HA-tagged S6K1-deltaCT), but not the wildtype (HA-tagged S6K1-WT), attenuated the JQ1-induced growth inhibition and apoptosis in COR-L279 cells. Two days after transfection, JQ1 was administered at the specified doses for 72 hours (b) or at 1 μM for 24 hours (c). Error bar represents SD. **(d)** Western blots show changes in BAD phosphorylation at S112, S136, and S155 after JQ1 treatment for 7 days at the specified doses. **(e)** Effects of RSK3 and S6K1-deltaCT overexpression on BAD phosphorylation two days after transfection in HEK293T cells. **(f)** A proposed model shows that BETi induces RSK3, which activates the TSC2-mTOR-p70S6K-BAD pathway to increase cell survival; mTOR inhibitor blocks this survival cascade and enhances BETi-induced apoptosis.

To identify the downstream effector of p70S6K1, we examined phosphorylation of the proapoptotic protein BAD, a known substrate of p70S6K1 (36). Phosphorylation of BAD releases antiapoptotic proteins BCL-2 and BCL-xl and promotes survival (37). After one-week treatment of JQ1, phosphorylation at BAD S112, but not at S136 and S155, increased in COR-L279 cells (Fig. 6d). A similar increase in BAD phosphorylation was observed after overexpression of RSK3 or S6K1-deltaCT in HEK293T cells, suggesting that RSK3 increases BAD phosphorylation via p70S6K1 (Fig. 6e).

We next assessed whether autophagy - a downstream pathway suppressed by mTOR, is involved in the synergy between mTORi and BETi. LC3B-II, a marker of autophagosome formation (38), was modestly induced by everolimus, consistent with autophagy induction after mTOR inhibition (Fig. S5). JQ1 did not induce LC3B-II, and neither did it affect everolimus-induced LC3B-II, suggesting that altered autophagy is not responsible for the synergy between mTORi and BETi (Fig. S5). Taken together, our findings support a model that RSK3 increases BAD phosphorylation via the TSC2-mTOR-p70S6K1 signaling cascade, and mTORi blocks this signaling and potentiates BETi-induced apoptosis (Fig. 6f).

## Discussion

Multiple investigational BETi drugs have been evaluated in early-phase clinical trials, and the results generally showed modest response rates and a short duration of response (39). Rational drug combinations could increase the antitumor efficacy of BETi and delay the development of drug resistance (39). In this study, we demonstrate that mTORi potentiates the antitumor effects of BETi in SCLC. RSK3 induction was identified as a resistance mechanism to BETi, and mTORi blocks the downstream signaling of this kinase and enhances BETi-induced apoptosis in SCLC.

Using high-throughput combination screens, we identified several classes of drugs synergizing with BETi in an unbiased manner. Some of these candidates, such as BCL2 inhibitors and HDAC inhibitors, have been shown previously to potentiate the antitumor effects of BETi in SCLC (7,10), while several others, such as inhibitors of cyclin-dependent kinases and carfilzomib, were reported to synergize with BETi in other tumor types (40,41). These earlier studies indirectly validate our results, suggesting shared vulnerabilities between various tumor types.

We focused on the combination of mTORi and BETi for further validation and mechanistic studies. Our results demonstrate that this combination has better antitumor effects than the single agents *in vitro* and *in vivo.* In addition, this combo was relatively well-tolerated, likely because the adverse effects of mTORi and BETi affect different organ systems. In contrast, the AKTi and BETi combo had poor tolerability *in vivo* despite being synergistic *in vitro*, probably because of overlapping adverse effects (e.g., diarrhea) of AKTi and BETi (42,43). These results stress the need to select the drugs with non-overlapping toxicities to increase antitumor activity and minimize adverse effects.

Unlike the findings in previous studies (13,15,16), our results did not support the feedback activation of RTKs after mTOR inhibition in SCLC. Previous studies have indicated epidermal growth factor receptor (EGFR) and insulin-like growth factor 1 receptor (IGF-1R) signaling pathways are commonly involved in the feedback activation of RTKs (14,16). In SCLC, EGFR signaling is inactive, while IGF-1R is upregulated in about 18.5% of the primary tumors (44). We suspect that the suppression of EGFR signaling accounts for the lack of feedback activation of RTKs following mTOR inhibition in SCLC.

Our study revealed *RPS6KA2* induction as a novel resistance mechanism to BETi in SCLC. The pro-survival function of RSK3 is mediated through mTOR signaling, which explains why mTORi would potentiate the antitumor effects of BETi. In breast and ovarian cancers, a higher expression of *RPS6KA2* has been associated with a worse prognosis, suggesting that this gene functions as an oncogene (45,46). Besides TSC2, RSK3 can also phosphorylate IκBα and causes activation of NF-κB (46). Previous studies in other neoplasms have reported that BETi resistance is caused by upregulation of WNT signaling via transcriptional program reset (47) or an increase of PI-3K-ERK activity through RTKs kinome rewiring (48). Our findings reveal a new player in the already diverse mechanisms of BETi resistance and confirm the central role of PI-3K-AKT-mTOR signaling activation.

In conclusion, we demonstrate that mTORi potentiates the antitumor effects of BETi in SCLC by blocking an RSK3-mediated survival signaling cascade. Our findings warrant further evaluation of mTORi and BETi combinations in SCLC patients.

## Authors’ Contributions

**A. Kumari:** Investigation, validation, methodology, writing–review and editing. **L. Gesumaria**: Investigation, validation, methodology, writing–review and editing. **Y. Liu**: Investigation, writing–review and editing. **VK Hughitt**: software, data curation, formal analysis, writing–review and editing. **X Zhang, M. Ceribelli, KM Wilson, C. Klump-Thomas, L. Chen, C. McKnight, Z. Itkin**: Investigation, software, data curation, formal analysis. **CJ Thomas:** Resources, formal analysis, funding acquisition, investigation, visualization, methodology, writing–review and editing. **BA Mock**: Conceptualization, funding acquisition, data curation, software, formal analysis, writing–review and editing. **DS Schrump**: Conceptualization, resources, funding acquisition, writing–review and editing. **H Chen**: Conceptualization, methodology, formal analysis, investigation, data curation, writing–original draft, writing–review and editing, visualization, supervision, project administration, funding acquisition.

## Acknowledgments

The authors thank Drs. Yulong Li, Shih-Hsin Hsiao, Kaitlin C. McLoughlin, Sichuan Xi, Ruihong Wang, Yuan Xu, Patricia Fetsch, and Markku M. Miettinen for their assistance in this study; Drs. Charles Rudin and John T. Poirier for providing the SCLC PDX models; Drs. Nenghui Wang and Mingzhu Yin for providing NHWD870.

## Supplemental Materials

**Table S2.** The results of the first high-throughput combination screens in H446 cells.

**Table S3.** The list of differential expression genes comparing the LX95 PDX tumors in the NHWD870, everolimus, or the combination groups to those in the vehicle group.

**Figure S1.**
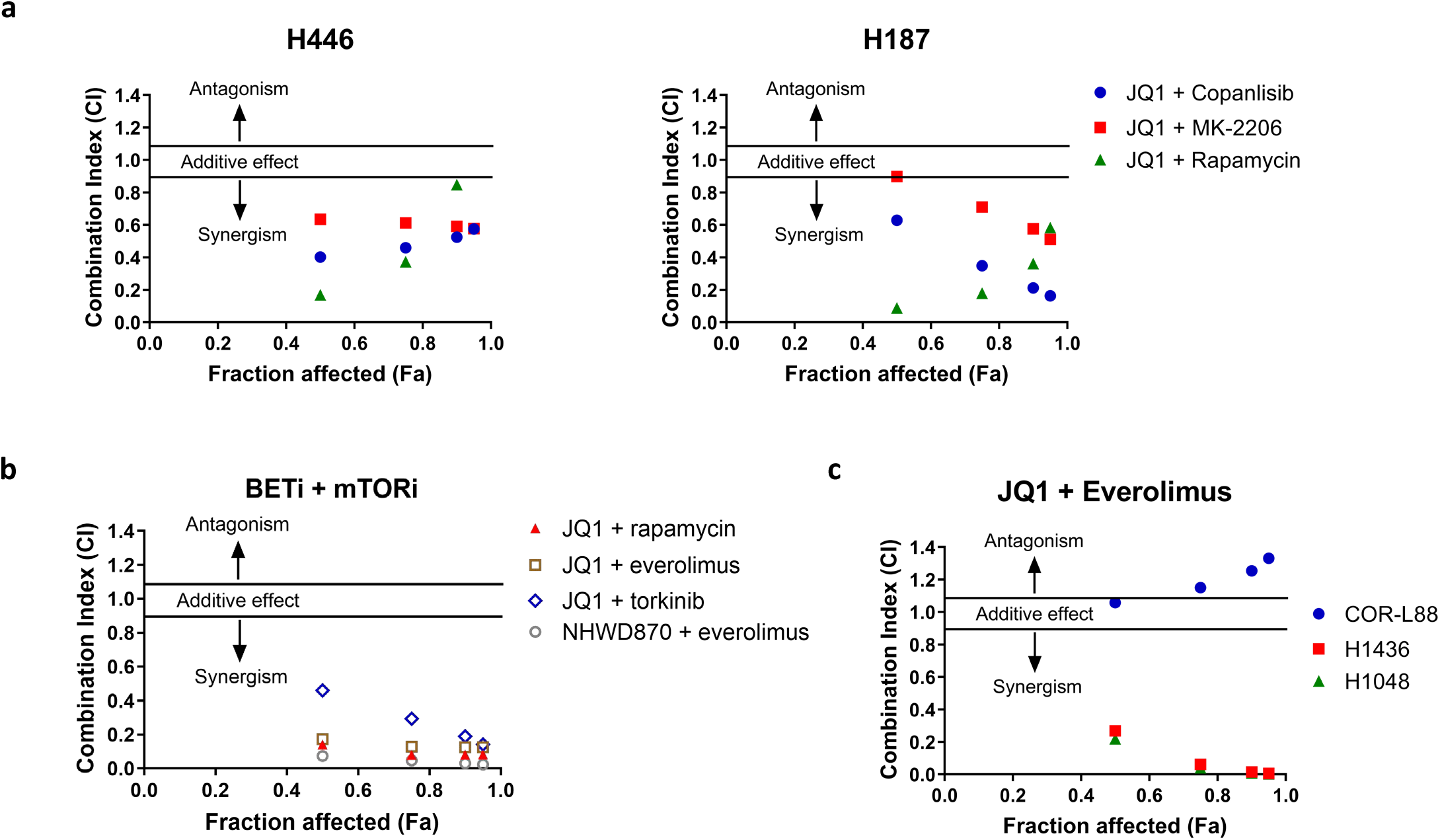
mTORi synergizes with BETi *in vitro*. **(a)** Synergy plots showing CIs versus affected fractions in H446 (left) or H187 cells (right) at 72 hours following treatment with JQ1 in combination with copanlisib, MK2206, or rapamycin. **(b)** Synergy plots showing CIs versus affected fractions in COR-L279 cells at 72 hours following treatment with various mTORi and BETi combinations as specified. **(c)** Synergy plot showing CIs versus affected fractions in three BETi-resistant SCLC lines (COR-L88, H1436, and H1048) at 72 hours following treatment with JQ1 in combination with everolimus.

**Figure S2.**
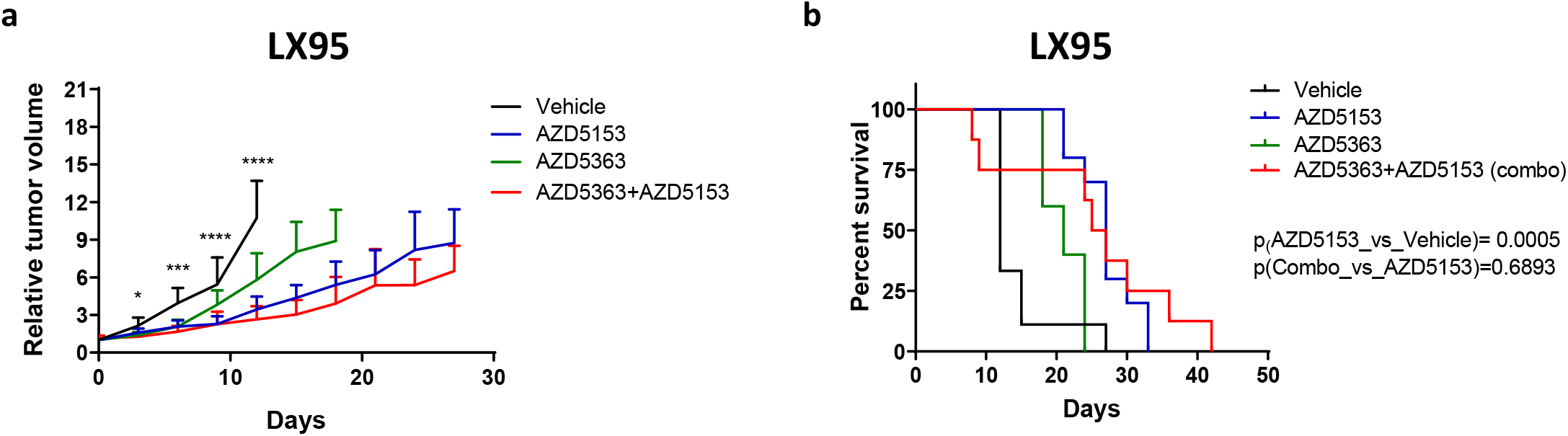
Compared to the single agents, the combination of AZD5363 (AKTi) and AZD5153 (BETi) failed to improve tumor control and survival in the LX95 PDX model. **(a)** Relative tumor volume changes in the first 29 days after drug administration. **(b)** Kaplan-Meier survival curves. LX95 PDX mice were initially treated with single-agent AZD5153 (1.5mg/kg, daily), single-agent AZD5363 (50mg/kg, daily), or the combination (AZD5153 1 mg/kg and AZD5363 50mg/kg, daily). Due to premature deaths in the combo group, the dose of AZD5363 in the combination group was reduced to 30mg/kg two weeks after the first dose. Black asterisks indicate statistically significant differences in tumor volumes between the AZD5153 single-agent group and Vehicle group, as determined by the Student’s *t*-test. No significant difference was found when comparing tumor volumes between the combo group and the AZD5153 single-agent group. The significance of the two-group comparisons in (b) was determined by the Log-rank test. Error bars represent SD. Where indicated, *, p<0.05; ***, p<0.001; ****, p<0.0001.

**Figure S3.**
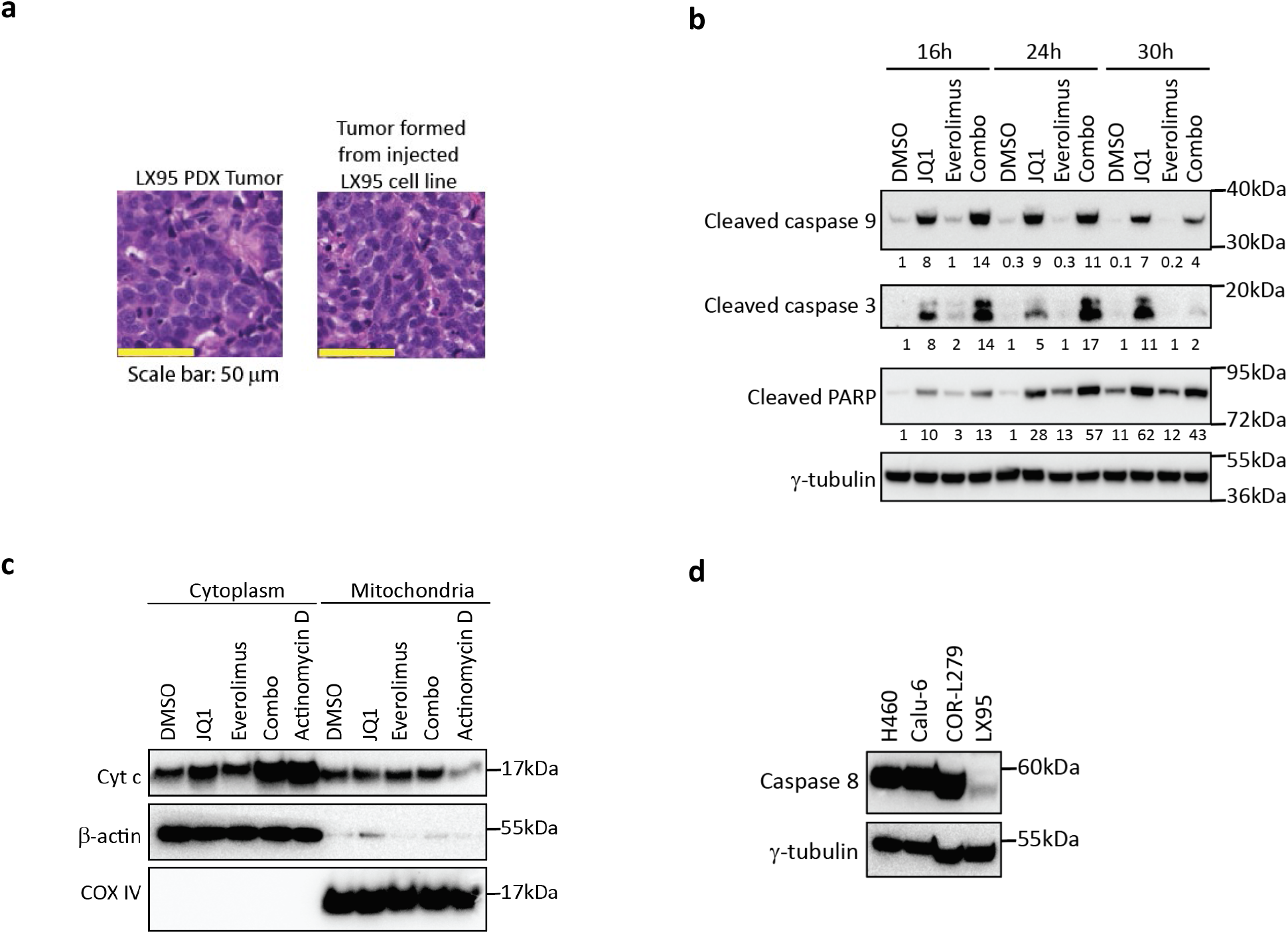
Everolimus enhances JQ1-induced apoptosis in LX95 cells via the intrinsic apoptotic cascade. **(a)** Hematoxylin and eosin staining of the xenograft tumors formed by an implanted LX95 PDX tumor block (left) or the LX95-derived cell line (right). **(b)** Western blots show cleavage of caspase 9, caspase 3, or PARP in LX95 cells at 16, 24, and 30 hours after treatment with JQ1 (156nM), everolimus (6.25nM), or the combination. **(c)** Western blots show cytochrome c abundance in the mitochondria and cytoplasmic fractions of LX95 cells at 16 hours after treatment with JQ1 (1μM), everolimus (6.25nM), or their combination. Actinomycin D (1μg/ml, 16 hours) was used as a positive control. **(d)** Western blots showing expression of caspase 8 in two non-small cell lung cancer lines (H460 and Calu-6) and two SCLC lines (COR-L279 and LX95).

**Figure S4.**
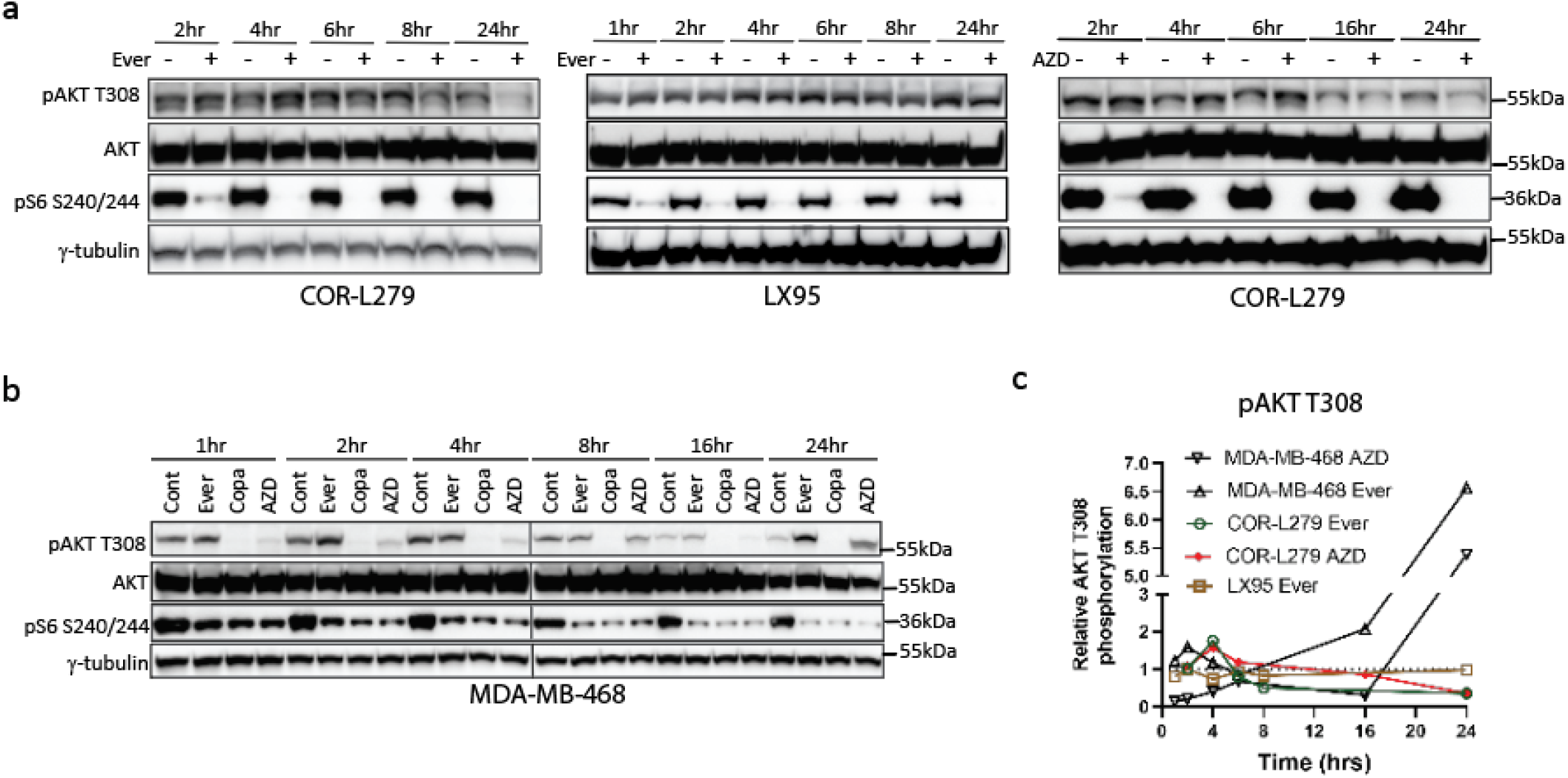
mTOR inhibition does not cause RTK feedback activation in SCLC. **(a)** Western blots show the dynamic changes of AKT T308 phosphorylation after everolimus treatment (5nM) in COR-L279 (left) or LX95 cells (middle), or after AZD8055 treatment (500nM) in COR-L279 cells (right). Phosphorylation of S6 S240/244 was examined to ensure mTOR inhibition after drug treatments. **(b)** Western blots show the dynamic changes of AKT T308 phosphorylation in MDA-MB-468 cells after treatment with everolimus (5nM), copanlisib (1μM), and AZD8055 (500nM). Copanlisib was used as a positive control. **(c)** Densitometry of AKT T308 phosphorylation in (a-b). AZD, AZD8055; Copa, copanlisib; Ever, everolimus.

**Figure S5.**
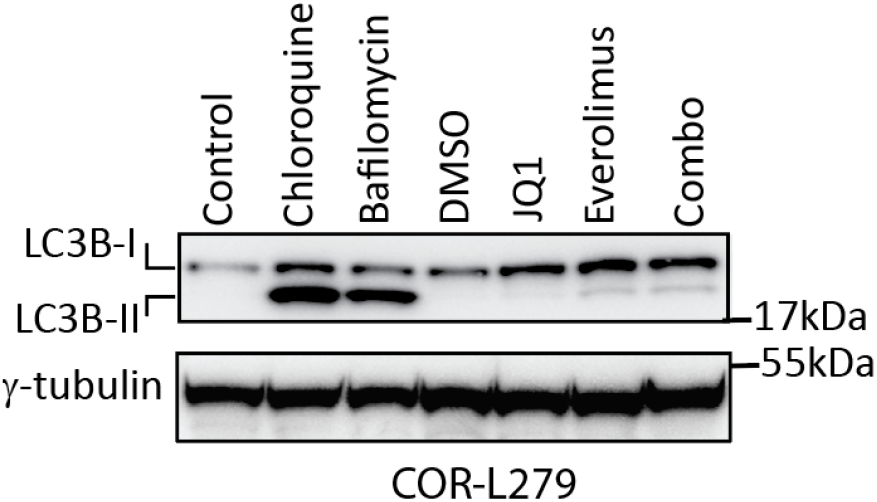
JQ1 does not affect everolimus-induced autophagy in COR-L279 cells. COR-L279 cells were treated with JQ1 (1μM), everolimus (6.25nM), or the combination for 24 hours before immunoblotting for LC3B-I and LC3B-II expression. Chloroquine (200 μM, 24 hours) and bafilomycin (150 nM, 24 hours) were used as positive controls.

**Table S1.**
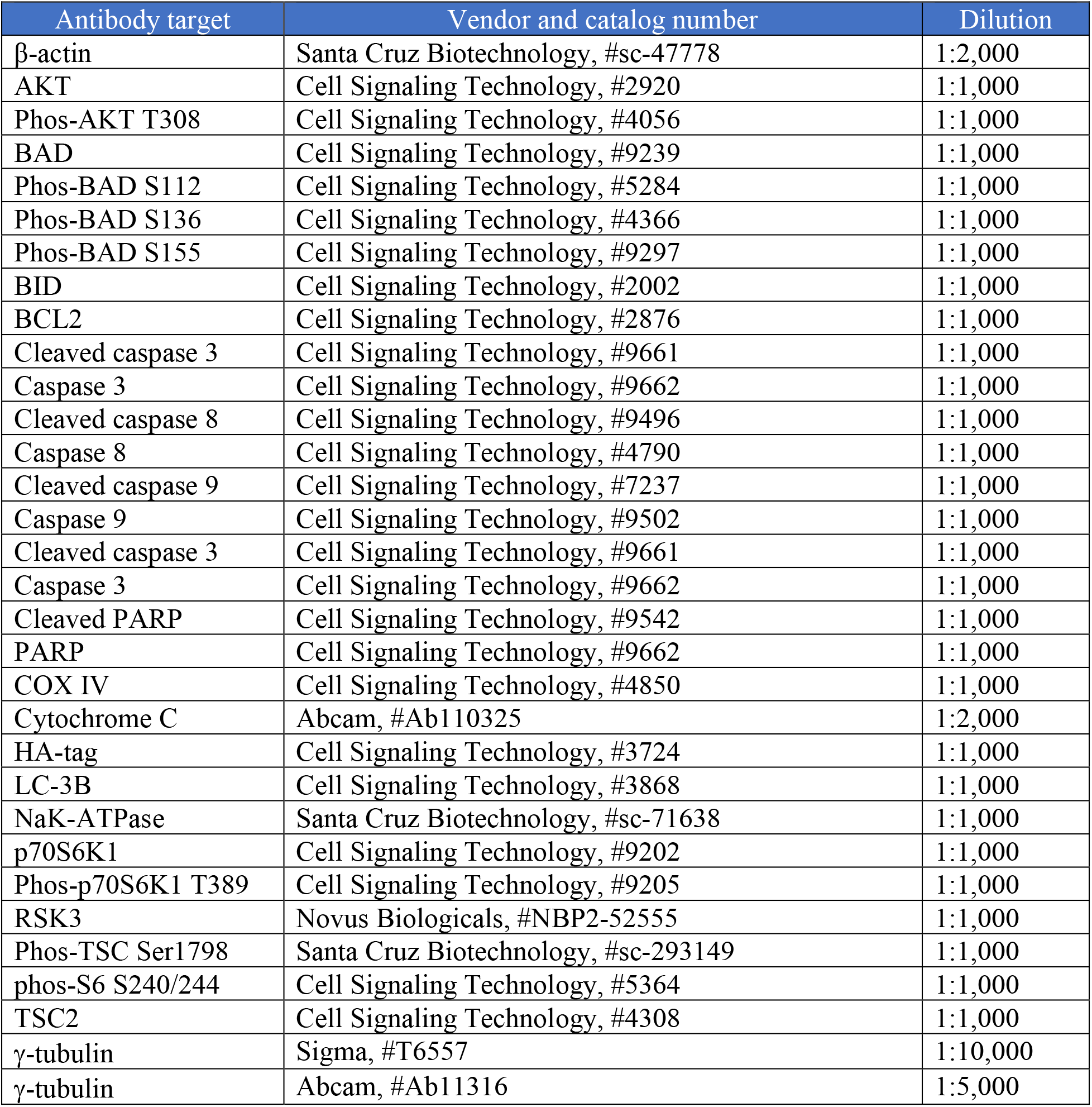
The list of the antibodies used in this study.

| Antibody target | Vendor and catalog number | Dilution |
| --- | --- | --- |
| β-actin | Santa Cruz Biotechnology, #sc-47778 | 1:2,000 |
| AKT | Cell Signaling Technology, #2920 | 1:1,000 |
| Phos-AKT T308 | Cell Signaling Technology, #4056 | 1:1,000 |
| BAD | Cell Signaling Technology, #9239 | 1:1,000 |
| Phos-BAD S112 | Cell Signaling Technology, #5284 | 1:1,000 |
| Phos-BAD S136 | Cell Signaling Technology, #4366 | 1:1,000 |
| Phos-BAD S155 | Cell Signaling Technology, #9297 | 1:1,000 |
| BID | Cell Signaling Technology, #2002 | 1:1,000 |
| BCL2 | Cell Signaling Technology, #2876 | 1:1,000 |
| Cleaved caspase 3 | Cell Signaling Technology, #9661 | 1:1,000 |
| Caspase 3 | Cell Signaling Technology, #9662 | 1:1,000 |
| Cleaved caspase 8 | Cell Signaling Technology, #9496 | 1:1,000 |
| Caspase 8 | Cell Signaling Technology, #4790 | 1:1,000 |
| Cleaved caspase 9 | Cell Signaling Technology, #7237 | 1:1,000 |
| Caspase 9 | Cell Signaling Technology, #9502 | 1:1,000 |
| Cleaved caspase 3 | Cell Signaling Technology, #9661 | 1:1,000 |
| Caspase 3 | Cell Signaling Technology, #9662 | 1:1,000 |
| Cleaved PARP | Cell Signaling Technology, #9542 | 1:1,000 |
| PARP | Cell Signaling Technology, #9662 | 1:1,000 |
| COX IV | Cell Signaling Technology, #4850 | 1:1,000 |
| Cytochrome C | Abcam, #Ab110325 | 1:2,000 |
| HA-tag | Cell Signaling Technology, #3724 | 1:1,000 |
| LC-3B | Cell Signaling Technology, #3868 | 1:1,000 |
| NaK-ATPase | Santa Cruz Biotechnology, #sc-71638 | 1:1,000 |
| p70S6K1 | Cell Signaling Technology, #9202 | 1:1,000 |
| Phos-p70S6K1 T389 | Cell Signaling Technology, #9205 | 1:1,000 |
| RSK3 | Novus Biologicals, #NBP2-52555 | 1:1,000 |
| Phos-TSC Ser1798 | Santa Cruz Biotechnology, #sc-293149 | 1:1,000 |
| phos-S6 S240/244 | Cell Signaling Technology, #5364 | 1:1,000 |
| TSC2 | Cell Signaling Technology, #4308 | 1:1,000 |
| γ-tubulin | Sigma, #T6557 | 1:10,000 |
| γ-tubulin | Abcam, #Ab11316 | 1:5,000 |

